# A Wireless Wearable Ecosystem for Social Network Analysis in Free-living Animals

**DOI:** 10.1101/2024.01.15.575769

**Authors:** Matt Gaidica, Mengxiao Zhang, Ben Dantzer

**Affiliations:** Department of Psychology, University of Michigan, Ann Arbor, MI, USA; School for Environment and Sustainability, University of Michigan, Ann Arbor, MI, USA; Department of Ecology & Evolutionary Biology, University of Michigan, Ann Arbor, MI, USA

**Keywords:** Wireless sensors, social networks, behavioral tracking, contact tracing, neuroethology

## Abstract

Understanding the dynamics of animal social systems requires studying variation in contact and interaction, which is influenced by environmental conditions, resource availability, and predation risk, among other factors. Traditional (direct) observational methods have limitations, but advancements in sensor technologies and data analytics provide unprecedented opportunities to study these complex systems in naturalistic environments. Proximity logging and tracking devices, capturing movement, temperature, and social interactions, offer non-invasive means to quantify behavior and develop empirical models of animal social networks. However, challenges remain in integrating different data types, incorporating more sensor modalities, and addressing logistical constraints. To address these gaps, we developed a wireless wearable sensor system with novel features (called “Juxta”), including modular battery packs, memory management for combining data types, reconfigurable deployment modes, and a smartphone app for data collection. We present data from a pilot study on prairie voles (*Microtus ochrogaster*), which is a small mammal species that exhibits relatively complex social behavior. We demonstrate the potential for Juxta to increase our understanding of the social networks and behavior of free-living animals. Additionally, we propose a framework to guide future research in merging temporal, spatial, and event-driven data. By leveraging wireless technology, battery efficiency, and smart sensing modalities, our wearable ecosystem offers a scalable solution for real-time, high-resolution data capture and analysis in animal social network studies, opening new avenues for exploring complex social dynamics across species and environments.

## I. INTRODUCTION

The dynamics of animal social systems are characterized by variations in contact and interaction that fluctuate both spatially and temporally. Specific drivers of variation in contact and interaction range from environmental conditions like weather, availability of resources, or the risk of predation. Yet, our understanding of these dynamics is often constrained by the limitations of traditional observational methods, which usually include direct observation of animals, especially in studies of free-living animals where focal animals are typically only observed for a snapshot in time or only specific behaviors are recorded. In this context, the advent of biological logging (sensor) technologies, animal-borne devices (“wearables”), and new analytical strategies has opened up unprecedented possibilities for studying the complex social behavior of animals, especially in naturalistic environments [1], [2].

Proximity logging and tracking devices are at the forefront of these technological advancements, offering novel opportunities to quantify movement, local temperature, and social interactions in a non-invasive manner [3]–[5]. These devices can capture high-resolution data that reveal subtle behavioral patterns and social structures, thereby enabling researchers to develop robust, empirical models of animal social networks. Techniques utilizing Bluetooth’s received signal strength indicator (RSSI, measured in dB) have become a useful means to approximate interaction distance between one-to-many individuals, something that is difficult to do with VHF or GPS [6]. The ability to collect such data is particularly valuable in the study of elusive, highly mobile, or nocturnal species, where conventional observational methods may prove inadequate or too disruptive [7].

Despite the significant progress in this field, several considerable challenges persist; the first pertains to integrating uniformly sampled temporal data (e.g., time-series data derived from accelerometers) with event-driven data (e.g., proximity detections). This challenge arises due to a disparity in the granularity of these data streams, which necessitates reconciliation. There is also a lack of standardized methodologies on how to represent these data in a probabilistic fashion, particularly when dealing with intricate, low-power sampling algorithms.

The second challenge is incorporating more sensor modalities into these studies to provide a richer context for social interactions. For instance, combining temperature and movement data in a wearable device could help estimate sleep-wake cycles given the data could be electrophysiologically validated [8]. Such multimodal data can enhance our understanding of how environmental features and internal states influence the social behavior and dynamics of animals. Lastly, current devices, while innovative and powerful, still present significant logistical challenges for use in free-living animals. These include constraints on battery life, weight and form, durability, and ease of deployment and retrieval. Specifically, in non-human animals, researchers are expected to adhere to the guideline that a wearable (animal-borne) device is no more than 5% of the body mass of an animal [9]. The ideal solution is multipurpose devices that integrate a range of sensor types into a single, lightweight unit, are easy to deploy on a wide variety of species and can be managed with portable tools like a smartphone.

Building upon the extensive research on the social behaviors of prairie voles, we sought to utilize our wireless wearable sensor technology, “Juxta”, to quantify multiple data streams (e.g., proximity detections, movement, collar temperature) while improving deployment ease through modular battery packs, rapidly printed and reusable collar cases, and a smartphone application for seamless data collection. Prairie voles, unlike most mammalian species, are known for their highly affiliative nature, forming enduring social bonds and displaying biparental behavior. These characteristics have made them an exciting candidate organism for studying social neuroscience [10]. With Juxta, we were able to capture high-resolution data on prairie vole social interactions in a free-living environment. Better understanding these dynamics could ultimately inform novel strategies for tackling psychiatric disorders associated with social cognitive deficits. Our pilot study represents a promising first step towards realizing this goal.

## II. FUNCTIONAL CONCEPT

### A. Electrical

Juxta devices are centered around a Texas Instruments CC2652P7 Bluetooth Low Energy (BLE) Microcontroller Unit (MCU; Fig 1a). This MCU is powered by a rechargeable 3.7V lithium polymer (LiPo) battery regulated to 1.8V using a high-efficiency dc-to-dc converter (Analog Devices ADP2108). The onboard voltage sensor was formed with a resistor divider.

**Fig. 1.**
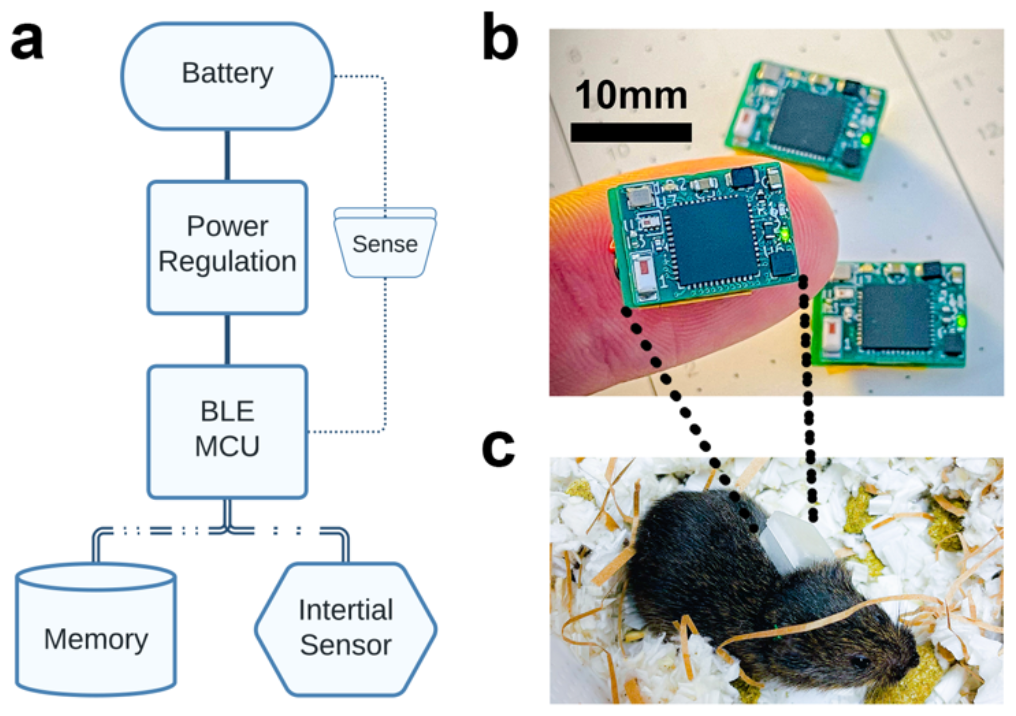
Juxta. (a) The battery is regulated to provide a stable power source to the Bluetooth Low Energy (BLE) Microcontroller Unit (MCU). A battery voltage sensor is implemented using two passive components. The memory and inertial sensor read/write over central communication lines. (b) The physical circuit board (c) packaged in a 3D-printed case.

The BLE MCU is connected via a standard serial peripheral interface (SPI) to 2Gb non-volatile NAND memory (Micron MT29F2G01) and an inertial sensor with temperature and magnetometer capability (STMicroelectronics LSM303AGR).

Other passive components include an impedance-matched balun and antenna (Johanson Technology 2450BM14G0011 and 2450AT18B100), battery/programming interface (Molex 505070-1022), a 48Mhz crystal oscillator, light emitting diodes, and capacitors.

### B. Physical

Juxta devices were fabricated using a 0.8mm FR4, 4-layer printed circuit board (PCB) process with physical dimensions of 11mm x 15mm (Fig. 1b). Modular battery PCBs were made separately with mating connectors (Molex 505066-1022) and batteries were hand-soldered. When mated with the battery (40mAh, procured from Alibaba), the entire functional device’s height was 8mm with a resulting weight of 3.2 grams; with the 3D-printed case and zip-tie, the total weight is 4.9 grams (Fig. 1c).

Custom battery charging boards were made to make charging easy and portable via USB by daisy-chaining four linear charge management controllers (Microchip MCP73831T).

Base station boards that interfaced with the devices served two purposes: enabling a large LiPo battery source to be input through a standard JST-PH connector and exposing a 10-pin JTAG connector (and other header pins) to program and debug devices

## II. SOFTWARE

### A. Embedded Software

The software running on each Juxta device was developed and compiled in Code Composer Studio and programmed using a Texas Instruments XDS 110 debugger. The devices included two main modes of operation: *shelf* and *interval*. In *shelf mode*, devices only advertise at a user-selected rate (typically, 1-10s), waiting for a connection. In *interval mode*, devices advertise but also scan (both user-selected rates). Interval mode also enabled periodic data logging of battery voltage, and device temperature via the inertial sensor. Data was stored in two separate areas of memory as either “log” data or “meta” data, described in detail below.

Log data were records consisting of a scanned device’s unique MAC address, RSSI, and time. The frequency of log data was ultimately determined by the scanning rate.

Meta data were records that were formatted as a data type, optional value, and time. Therefore, meta data were general-purpose records used for both behavioral/sensor data as well as key events useful for debugging and analysis. The events consisted of an accelerometer event (movement exceeded a user-defined threshold), battery voltage (an example of a record that includes data), collar temperature, the current device mode, as well as user-initiated events such as establishing a connection or resetting the memory.

Log and meta data are accessed and exported separately (see Smartphone Application).

Devices automatically detected if they were deployed as a base station or on an animal. Base stations increased their radio power for further-ranging advertisements and sleep modes were disabled to increase clock accuracy. Therefore, base stations were meant to act as central time coordinators. Animal-based devices exploited a custom field in their advertisement that would flag base stations for a time update (set to once every day). If this flag was detected by a base station during a scan, it would attempt to form a connection and synchronize time.

The embedded software exposed individual BLE characteristics that act as the interface between the device and the outside world; they are readable (R) or writable (W). In brief, these characteristics were for time (R/W), log and meta record counts (R), mode (R/W), subject (R/W), voltage (R), and collar temperature (R). There were also two characteristics for data transfer: one to accept commands (e.g., dump log data) and the other to stream data packets using BLE notifications.

### B. Smartphone Application

The smartphone application was built using XCode and deployed on iOS devices (e.g., Apple iPhone) using TestFlight (Apple Inc.). The application only scanned and connected to devices that had the BLE characteristics previously described. Upon connecting to a device, the user was presented with an overview of the current settings, including mode, subject, sync/reset, and the ability to dump log or meta records (Fig. 2a). The Advanced Options pane enabled users to modify the advertise and scan intervals (Fig. 2b).

**Fig. 2.**
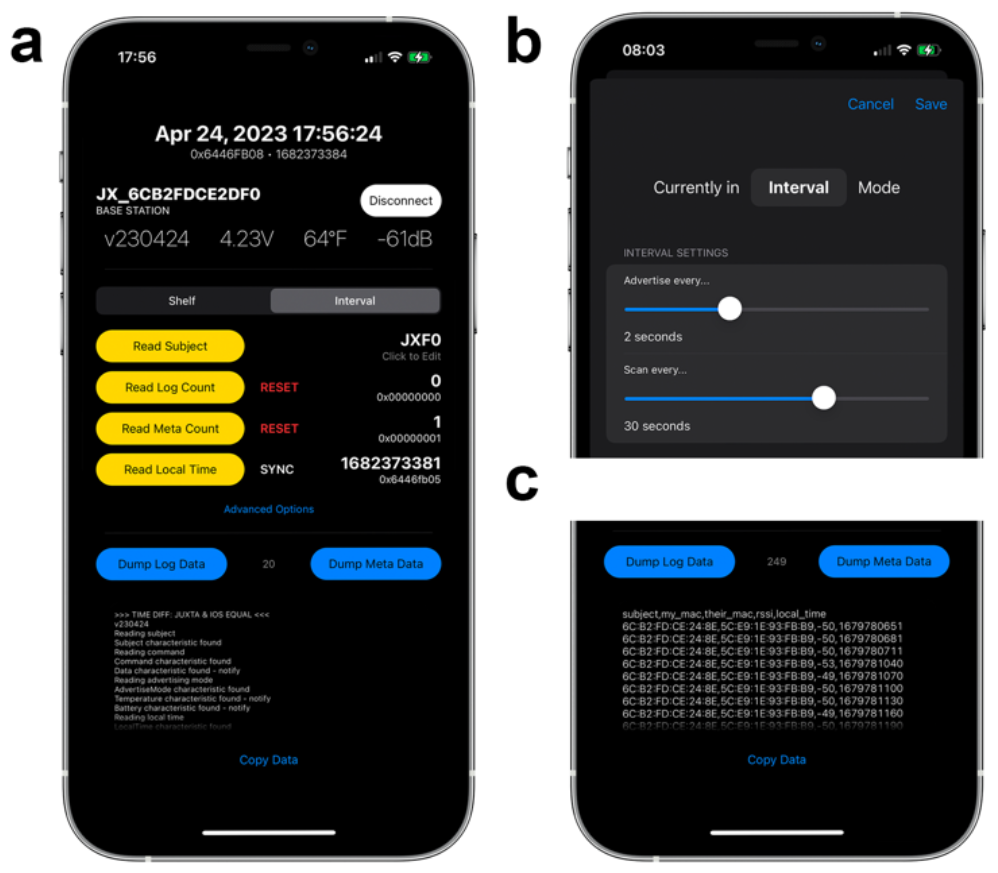
Smartphone Application. (a) The main interface connected to a device. The upper third shows device information, the middle third read/writes device settings, and the lower third contains a data terminal. (b) The advanced Options panel. (c) Terminal output for log data.

When a user initiated data dumping (either log or meta data), the data appeared as rows in the main textbox (Fig. 2c). The app continues to count those rows indicating the number of records retrieved. When complete, the comma-delimited data from the textbox can be copied to the phone’s clipboard and pasted into a data repository.

## IV. DEPLOYMENT

The devices were protected by a custom two-piece enclosure 3D-printed (Anycubic Photon Mono 4K) using an ABS-like resin. All-purpose RTV silicone (JB Weld) was applied to the enclosure interface to weatherproof devices, after which they were secured as collars to free-ranging voles using zip ties (Uxcell, 2mm x 150mm) with heat shrink tubing applied over the locking mechanism to avoid it from shifting.

Our pilot data originated from 96 collared prairie voles (48 females, 48 males) released into four enclosures of 0.1 hectares at the Miami University Ecology Research Center (ERC; Oxford, Ohio). Every 4 days, attempts were made to recapture voles and change batteries. Details of the capture and animal handling procedures can be found elsewhere [11], [12]. During the recapture event, data was retrieved from the devices and archived.

The average lifetime of a device in shelf mode is over 30 days, allowing devices to be prepped ahead of time. When set to interval mode for deployment, we found a good tradeoff of advertising every 10s and scanning every 60s, leading to a lifetime of 6-7 days.

## V. ANALYSIS

Log and meta data were combined in MATLAB to generate analytics. Time-correlated data enable several insights including approximate periods of wakefulness (e.g., when activity goes down and the temperature goes up; Fig. 3a) as well as the time course of interactions (Fig. 3b).

**Fig. 3.**
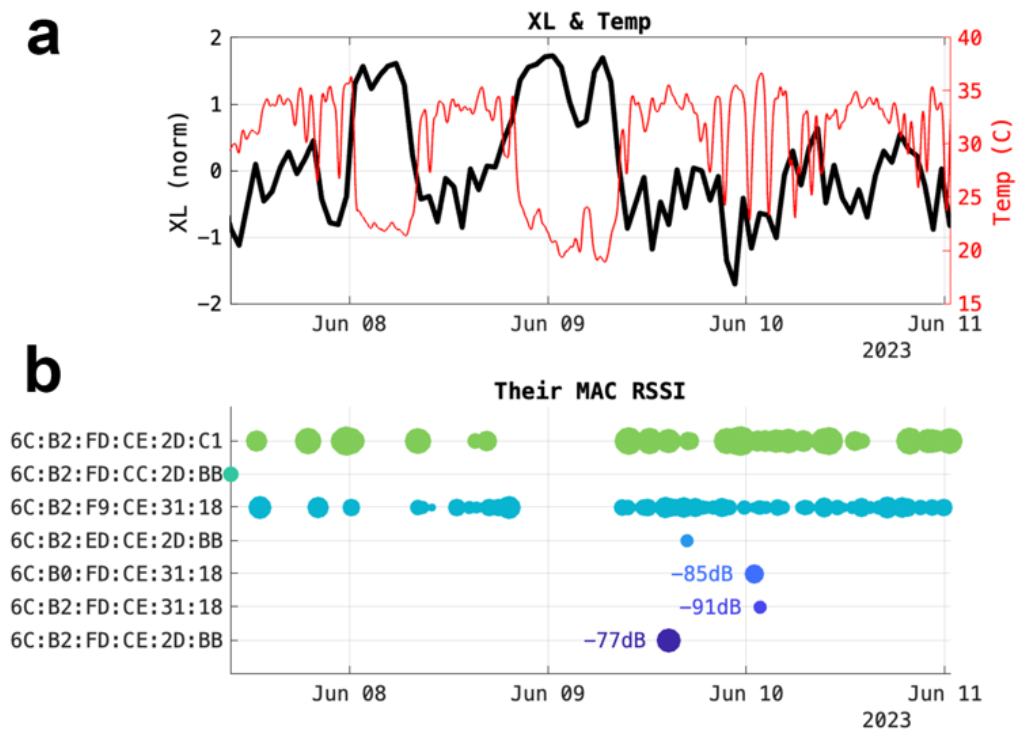
Exemplar Data. (a) Inertial threshold events (XL) and temperature data (Temp) over time. (b) Scanning activity of unique (colored) nearby devices over time. The y-axis contains the other device’s MAC address (Their MAC) and the size of the circle represents the RSSI, a proxy for physical distance. Three values are labeled to indicate how circle size correlates to RSSI.

To determine the correlation between RSSI and animal-animal distance we ran a manual, indoor test with 5 devices set known distances apart. We observed a typical RSSI fall-off with the inverse square of distance where an RSSI of greater than -80dB meant the collars were within 5cm of each other with a max detection distance of roughly 20cm (equating to an RSSI of less than -95dB).

These data can be easily transformed to generate insights related to circadian rhythms [13] or standard social network strength between one-to-many individuals [14].

## VI. CONCLUSION

In this paper, we delve deeper into the novel features of our wireless wearable sensor system, Juxta, elucidating how each of its unique characteristics—ranging from the modular battery packs to the versatile data collection capabilities—addresses the existing challenges in studying free-living animal social networks, thereby enriching the depth and breadth of data acquisition in this field [1]. Central to the challenge of merging temporal, spatial, and event-driven data, we propose a framework that could guide future interdisciplinary research.

The primary engineering constraint for future developments will continue to rest between physical size and optimization of power— which is highly dependent on sampling modes—to provide rich data and facilitate long-duration field deployments. This is true for wearable and implantable devices, and it should be noted that implantables have the potential added benefit of being shielded from very cold ambient temperatures that reduce battery life. But despite increasingly smaller and more efficient Bluetooth Low Energy (BLE) Microcontroller Units (MCUs), battery technology has not advanced as rapidly. Although rechargeable batteries make sense from a reuse standpoint, their power density is roughly an order of magnitude less than their non-rechargeable counterparts.

We propose exploring alternative proximity detection technologies to enhance power efficiency. For instance, intimate interactions may leverage extremely low-power ferromagnetic or capacitive sensors positioned around a collar to cue the radio to wake up instead of relying on less efficient, fixed sampling routines. This would require rethinking the device form factor and potentially using novel flexible PCB substrates. Likewise, more intelligent radio modes could be implemented. Perhaps interactions are most common and important when inertial activity is detected or highly correlated with the circadian phase, in which case devices could emphasize detections during those periods. While it could be argued that irregular or machine-trained sampling modes complicate interpretation and analysis, those barriers sit squarely between the past and future approaches to quantifying behavior and interactions in necessarily constrained environments and model systems (e.g., the prairie vole).

Whether BLE and the 2.4GHz wireless band are the most suitable protocol and technology for this application remains debatable. Our study examined the capability to monitor interactions of a tunneling, terrestrial mammal. Extending the use case to larger territories or 3D environments presents several challenges. Firstly, there is a tradeoff between signal power (or amplification) and device lifespan. Secondly, accommodating an antenna in a small space result in anisotropic radiation patterns, complicating signal strength interpretation. Compared to more efficient and far-reaching wireless protocols (e.g., Sub-1 GHz technologies), it becomes difficult to reconcile the use of BLE with the advantages of alternatives [15]. However, the prevalent use of BLE in wearable consumer electronics has expedited device development, miniaturization, advancements in distance estimates and positional accuracy, and support across mobile platforms. Therefore, BLE shortcomings might be outweighed by its future potential and ease of implementation [5].

By leveraging advances in wireless technology, battery efficiency, and smart sensing modalities, our wearable ecosystem offers a scalable, flexible solution for real-time, high-resolution data capture and analysis in animal social network studies. We are actively merging this technology into implantable form factors to record neural and cardiac biorhythms alongside social contacts for both laboratory and field experiments [8]. Together, what we present opens new or tangential avenues for the exploration of complex social dynamics and other emerging questions in a wide range of species and environments.

## ACKNOWLEDGMENT

We would like to thank ERC Director Ann Rypstra and ERC Manager Jeremy Fruth. We thank James Burkett (University of Toledo) for providing the prairie voles used in this research. This work was directly supported by NIH Project 5R21MH127500-02 and by the Translational Research Institute for Space Health through NASA Cooperative Agreement NNX16AO69A.

